# Proteomic analysis of microtubule inner proteins (MIPs) in Rib72 null *Tetrahymena* cells reveals functional MIPs

**DOI:** 10.1101/2020.10.02.324467

**Authors:** Amy S. Fabritius, Brian A. Bayless, Sam Li, Daniel Stoddard, Westley Heydeck, Christopher C. Ebmeier, Lauren Anderson, Tess Gunnels, Chidambaram Nachiappan, Justen B. Whittall, William Old, David A. Agard, Daniela Nicastro, Mark Winey

## Abstract

Motile cilia and flagella are built from stable populations of doublet microtubules that comprise their axonemes. Their unique stability is brought about, at least in part, by a network of <u>M</u>icrotubule <u>I</u>nner <u>P</u>roteins (MIPs) found in the lumen of their doublet microtubules. Rib72A and Rib72B were identified as microtubule inner proteins (MIPs) in the motile cilia of *Tetrahymena thermophila*. Loss of these proteins leads to ciliary defects and loss of multiple MIPs. We performed mass spectrometry coupled with proteomic analysis and bioinformatics to identify the MIPs lost in *RIB72A/B* knockout (KO) *Tetrahymena* cells. From this analysis we identified a number of candidate MIPs and pursued one, Fap115, for functional characterization. We find that loss of Fap115 results in disrupted cell swimming and aberrant ciliary beating. Cryo-electron tomography reveals that Fap115 localizes to MIP6a in the A-tubule of the doublet microtubules. Overall, our results highlight the complex relationship between MIPs, ciliary structure, and ciliary function.

## Introduction

Microtubules perform diverse essential functions within the cell, such as acting as tracks for movement of cargos, forming the mitotic spindle, and serving as the major structural constituent of cilia/flagella. Functional differentiation of microtubules depends heavily on combinations of different post-translational modifications as well as microtubule associated proteins (MAPs) that bind the outside of microtubules (Bodakuntla et al., 2019; Janke, 2014).

*In vitro*, microtubules display tremendous dynamicity. The ability of microtubules to rapidly grow and shrink is essential for a number of their functions. However, not all populations of microtubules are dynamic. Microtubules found in the axons of neurons and the axonemes of cilia and flagella do not display dynamic properties. In the case of motile cilia/flagella axonemes, the microtubules are also resistant to the bending forces that cytoplasmic microtubules are susceptible to (Portran et al., 2017; Xu et al., 2017). This stability is critical for motile cilia/flagella function because these microtubules must bend to create a persistent beating motion that moves extracellular fluid. In humans, an inability of motile cilia to generate extracellular fluid flow results in a wide range of maladies including hydrocephaly, respiratory difficulty, and female infertility (Lyons et al., 2006; Sawamoto et al., 2006; Wanner et al., 1996). Gaining an understanding of how motile cilia/flagella are assembled and stabilized is critical to understanding the pathology of these poorly characterized disorders.

The structure of motile cilia/flagella axoneme microtubules may be responsible for their unique stability. Unlike cytoplasmic microtubules, ciliary/flagellar axoneme microtubules are always found as a doublet of two microtubules. They consist of a single 13 protofilament A-tubule joined to a 10 protofilament B-tubule that shares a wall with the complete A-tubule (Fig. 1A). Microtubules do not naturally form doublets *in vitro*, which suggests that other genetic determinants are necessary to create this modified microtubule structure.

**Figure 1:**
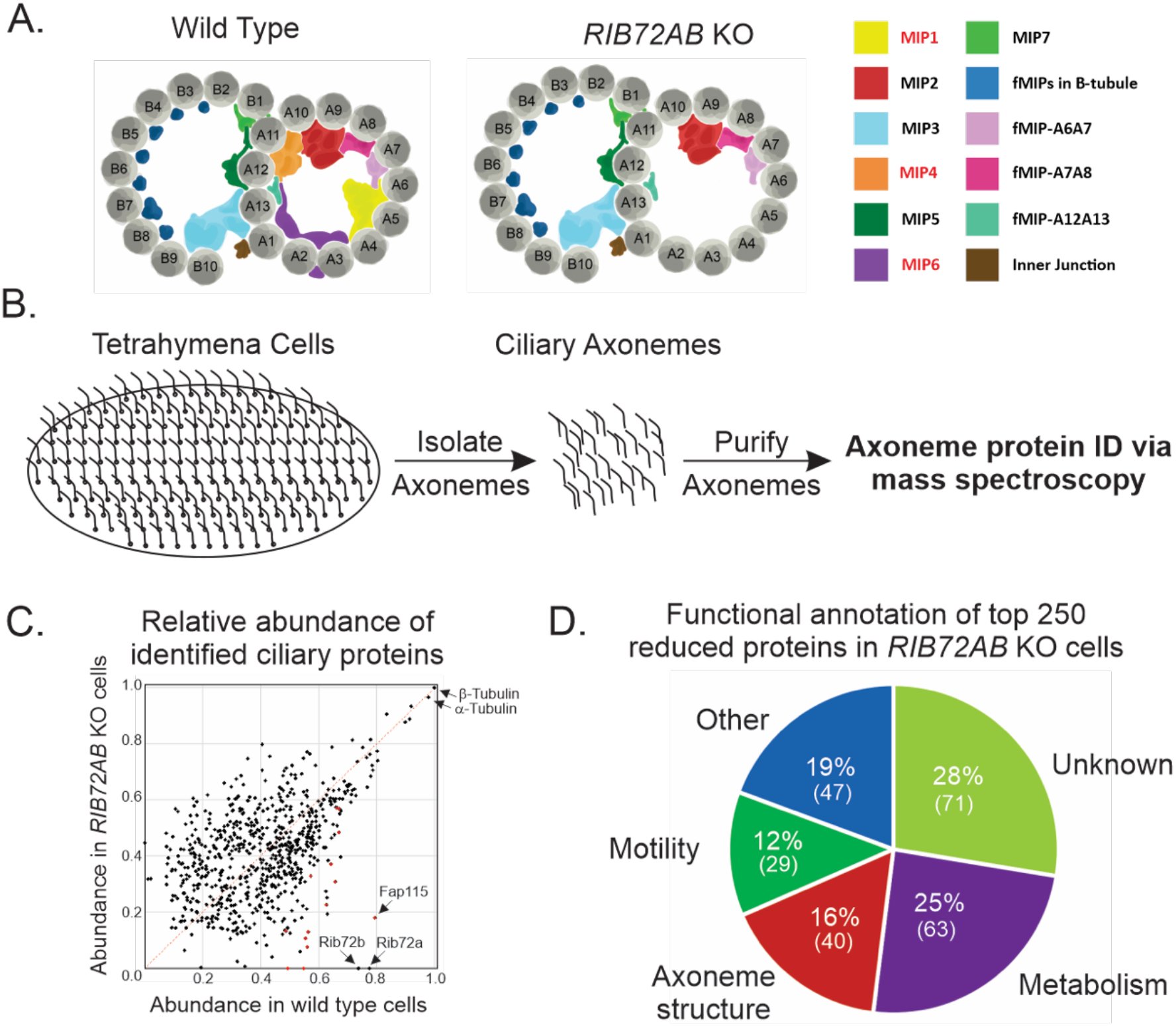
Identification of proteins lost in *RIB72AB* knockout *Tetrahymena* cells. A) Schematic representation of ciliary doublet microtubules in wild type (left) and *RIB72AB* KO cells showing the loss of A-tubule MIPs. B) Workflow for the identification of proteins lost in *RIB72AB* KO doublet microtubules. Axonemes were isolated from wild type, *RIB72A* KO, *RIB72B* KO, and *RIB72AB* KO cells and analyzed by mass spectrometry. C) Scatterplot showing normalized abundance of doublet microtubule proteins in wild type vs *RIB72AB* KO mutant cells. Black dots represent proteins and red dots represent candidate MIP proteins. All proteins that fall under the red dashed line are reduced in *RIB72AB* KO cells relative to wild type cells. D) Functional annotation of top 250 proteins lost in *RIB72AB* KO cells based on domain analysis and known homolog characterization

Single particle cryo-electron microscopy (Cryo-EM) and cryo-electron tomography (Cryo-ET) have led to the exciting discovery of a network of non-tubulin densities within the lumen of doublet microtubules (Ichikawa and Bui, 2018). These densities have been termed <u>M</u>icrotubule <u>I</u>nner <u>P</u>roteins, or MIPs. Several recent studies have identified the proteins and potential functions of some MIPs in ciliates and flagellates (Ichikawa et al., 2017; Khalifa et al., 2020; Kirima and Oiwa, 2018; Ma et al., 2019; Owa et al., 2019; Stoddard et al., 2018; Urbanska et al., 2018). High-resolution structures, especially in *Chlamydomonas reinhardtii* doublet microtubules, provide a strong basis for identification and characterization of these proteins (Ma et al., 2019). While some studies have provided clues to the functional significance, e.g. microtubule lattice spacing and angles and intolerance to mechanical stress (Ichikawa et al., 2017, 2019; Owa et al., 2019), there is still a need for additional information about the roles of individual MIPs and MIPs as a group. In addition, some MIP proteins have yet to be identified and characterized, and MIPs are likely to vary among species and systems.

We previously characterized the MIPs Rib72A and Rib72B in the ciliate *Tetrahymena thermophila* (Stoddard et al., 2018). We found that Rib72 is necessary for the localization of a majority of the MIPs found in the A-tubule of doublet microtubules and that loss of Rib72 proteins results in reduced ciliary beating function and doublet microtubule stability. Here, we report our proteomics- and bioinformatics-based approach to identify the MIPs lost in *RIB72*KO *Tetrahymena* cells. We follow this up with functional characterization of a single MIP candidate, Fap115. Through the use of Cryo-ET we identify that Fap115 is a component of MIP6a, a previously identified structure in the A-tubule of doublet microtubules (Stoddard et al., 2018). Furthermore, we find that the power stroke of motile cilia beating is slowed in *FAP115* KO cells. Overall, our work not only helps us begin to understand the complex network of MIPs within doublet microtubules but importantly it demonstrates an experimental pipeline that will allow for the functional characterization of other candidate MIPs. These steps are essential for understanding the molecular mechanism of cilia assembly.

## Results and Discussion

### Rib72 phylogeny

Our previous studies identified *Tetrahymena* Rib72A and Rib72B as MIPs and showed that they are necessary for the localization of a majority of A-tubule MIPs (Fig. 1A) (Stoddard et al., 2018). To better understand whether Rib72A and Rib72B (and A-tubule MIPs) are essential for the formation of motile cilia, we queried a wide panel of eukaryotes (both containing and not containing motile cilia) for the presence of Rib72 homologs (Table 1). We find that a majority (15 of 21) of species that have motile cilia also have a homolog for Rib72. Almost every species that does not contain motile cilia correspondingly does not have Rib72. One exception is *C. elegans*. Interestingly, while *C. elegans* does not contain motile cilia, they do contain sensory cilia, suggesting that sensory cilia may contain MIPs (Inglis et al., 2018). Given the non-overlapping roles of Rib72A and Rib72B in *Tetrahymena* we also wanted to assess whether other species had two Rib72 proteins. We note that the majority of motile cilia containing species that have Rib72A (9 of 14) do not have a homolog for *Tetrahymena* Rib72B, which suggests that Rib72A is sufficient to recruit all Rib720dependent A-tubule MIPs in species without Rib72B. Overall, our findings highlight the importance of Rib72 in the formation of motile cilia and its absense in organisms not containing motile cilia.

**Table 1:**
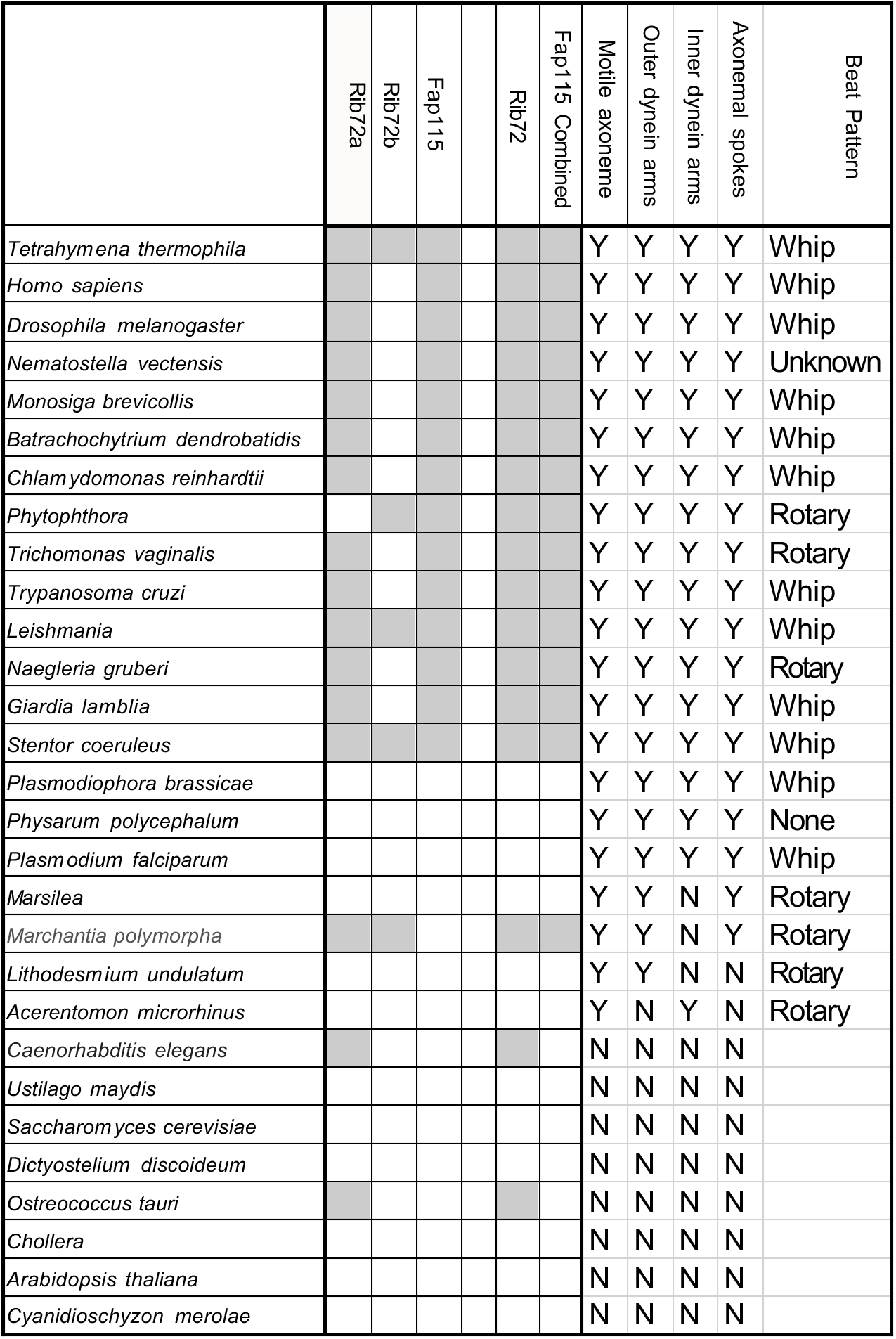
Homology assessment based on best-hit reciprocal BLAST

### Proteomics/Analysis

Determining the molecular identity of MIPs is essential for our understanding of cilia structure. Since the loss of Rib72A and Rib72B in *Tetrahymena* leads to loss of a majority of MIP densities within the A-tubule (Fig. 1A), we sought to identify these missing MIPs by a proteomics-based approach. To do this, we compared relative protein abundance between cilia isolated from wild-type and *RIB72A/B* double KO *Tetrahymena* cells (Fig. 1B). Mass spectrometry analysis of *RIB72A/B* KO protein levels revealed ~2100 proteins with altered abundance compared with the wild-type protein levels. From the 2100 proteins, the top 1000 most abundant proteins in wild-type cells were plotted based on their relative abundance in wild-type axonemes vs *RIB72A/B* KO axonemes (Fig. 1C). The top 250 proteins from this list were annotated for predicted functional role (Fig. 1D). There were 40 proteins that were found to associate with microtubules or have homologs that function at the cilia/flagellar axoneme. They were considered potential structural proteins of the cilia (Fig. 1D). This list is not fully inclusive and there is potential that some candidate MIPs were not included, so in addition to these potential axoneme structural proteins we also looked at the top 50 proteins that are lost in *RIB72A/B* KO cells, regardless of functional classification (Table S1). To these proteins, we applied various bioinformatics approaches, including reciprocal besthit BLAST analysis, domain analysis, comparisons with previously published basal body and ciliary proteomes, and findings from the recent *Chlamydomonas* MIP map (Keller et al., 2005; Kilburn et al., 2007; Ma et al., 2019) to remove proteins that are unlikely to be *Tetrahymena* MIPs and to narrow the list to a reasonable number for analysis. The identity of the candidates based on available information is shown in Table 2.

**Table 2:**
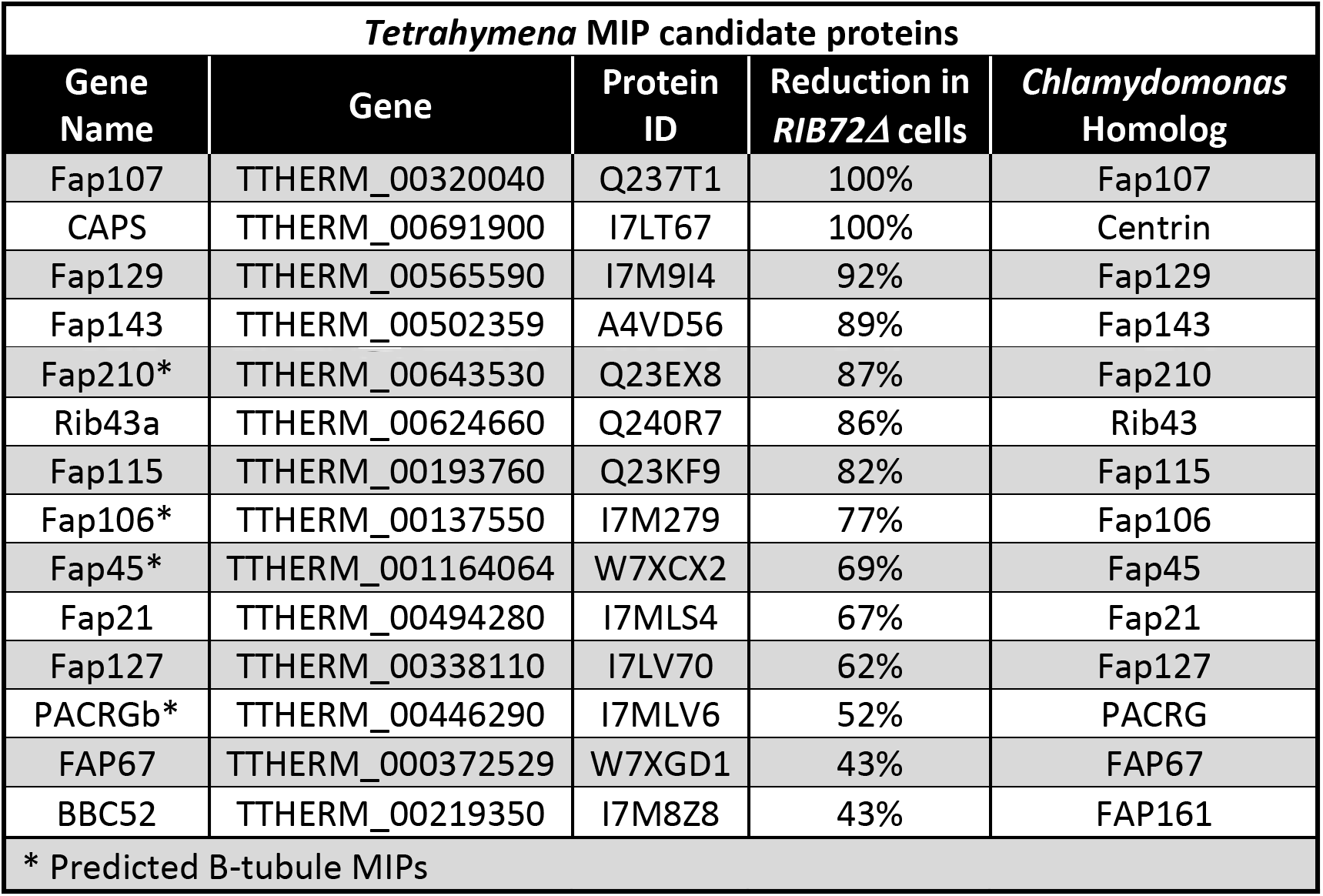
Candidate Tetrahymena MIP proteins

| ***Tetrahymena* MIP candidate proteins** |  |  |  |  |
| --- | --- | --- | --- | --- |
| **Gene Name** | **Gene** | **Protein ID** | **Reduction in *RIB72Δ* cells** | ***Chlamydomonas* Homolog** |
| Fap107 | TTHERM_00320040 | Q237T1 | 100% | Fap107 |
| CAPS | TTHERM_00691900 | I7LT67 | 100% | Centrin |
| Fap129 | TTHERM_00565590 | I7M9I4 | 92% | Fap129 |
| Fap143 | TTHERM_00502359 | A4VD56 | 89% | Fap143 |
| Fap210* | TTHERM_00643530 | Q23EX8 | 87% | Fap210 |
| Rib43a | TTHERM_00624660 | Q240R7 | 86% | Rib43 |
| Fap115 | TTHERM_00193760 | Q23KF9 | 82% | Fap115 |
| Fap106* | TTHERM_00137550 | I7M279 | 77% | Fap106 |
| Fap45* | TTHERM_001164064 | W7XCX2 | 69% | Fap45 |
| Fap21 | TTHERM_00494280 | I7MLS4 | 67% | Fap21 |
| Fap127 | TTHERM_00338110 | I7LV70 | 62% | Fap127 |
| PACRGb* | TTHERM_00446290 | I7MLV6 | 52% | PACRG |
| FAP67 | TTHERM_000372529 | W7XGD1 | 43% | FAP67 |
| BBC52 | TTHERM_00219350 | I7M8Z8 | 43% | FAP161 |
| * Predicted B-tubule MIPs |  |  |  |  |

In addition to other A-tubule MIPs, loss of Rib72 may affect the localization of B-tubule MIPs. In our proteomic analysis we find a number of B-tubule MIPs to be reduced in *RIB72A/B* KO *Tetrahymena* cells. This raises the intriguing possibility that the assembly of the A-tubule directly impacts the formation of the B-tubule. Some MIPs have been shown to reach across the tubulin lattice and it is possible that these projections into the B-tubule could serve as a platform for B-tubule specific MIP localization (Ichikawa et al., 2017) (Ma et al., 2019). A-tubule MIPs also make connections with dynein arms and radial spoke proteins (Ichikawa et al., 2019; Khalifa et al., 2020). The propagation effect of lost connections from the inside of the axoneme to the accessory structures that are necessary for beating and ciliary waveform can explain the ciliary beating defects associated with MIP loss. Overall, the intricacies of MIP interaction across the A- and B-tubule, along with forging connections to dyneins and radial spoke proteins, suggests a high level of interdependence among all structural components of axonemes.

One of the MIP candidates, Fap115, was of particular interest to us because it shows an 82% reduction in abundance in *RIB72* double KO cilia relative to wild-type cilia and was previously identified as a component of the *Tetrahymena* basal body (Kilburn et al., 2007). In addition, *Chlamydomonas* Fap115 is present inside the A tubule of the axoneme (Ma et al., 2019). We also find that 100% of motile cilia containing species that contain Rib72 also contain Fap115 (Table 1), suggesting potential essential interaction or Rib72-dependent function. Overall, our proteomics approach was successful in generating a list of candidate *Tetrahymena* MIPs, of which Fap115 was chosen as the single protein of most immediate interest.

### Fap115 is a ciliary protein that is necessary for ciliary beating

Fap115 in *Tetrahymena* is a ~110 kDa, EF-hand domain containing protein and was one of the original MIP candidates found in earlier 2D gel screens of salt-extracted axonemes of *Chlamydomonas* flagella in the Nicastro lab (data not shown). Recently, a new and extensive *Chlamydomonas* mutant library was generated (Li et al., 2016), containing a fap115 mutant. After PCR verification of the mutant, we performed cryo-ET analysis. Isolated axoneme cryo-ET revealed a single defect in the MIP structure, that of MIP6a along PFA2. MIP6a was missing, whereas all other MIP subunits were present within the subtomogram averages (Figure S1). This is consistent with the localization of fap115 in *Chlamydomonas* by Ma et al. (2019).

To further confirm the role and identity of Fap115 as a MIP, we analyzed its function in *Tetrahymena* cilia. We first confirmed the localization of Fap115 to *Tetrahymena* cilia and basal bodies by creating an mCherry C-terminal fusion (Fig. 2A). We then created a *Tetrahymena* knockout strain of *FAP115*, which was confirmed by RT-PCR. The FAP115 KO strain was used to analyze the potential role of Fap115 as a MIP in stability or function of the ciliary axoneme.

**Figure 2.**
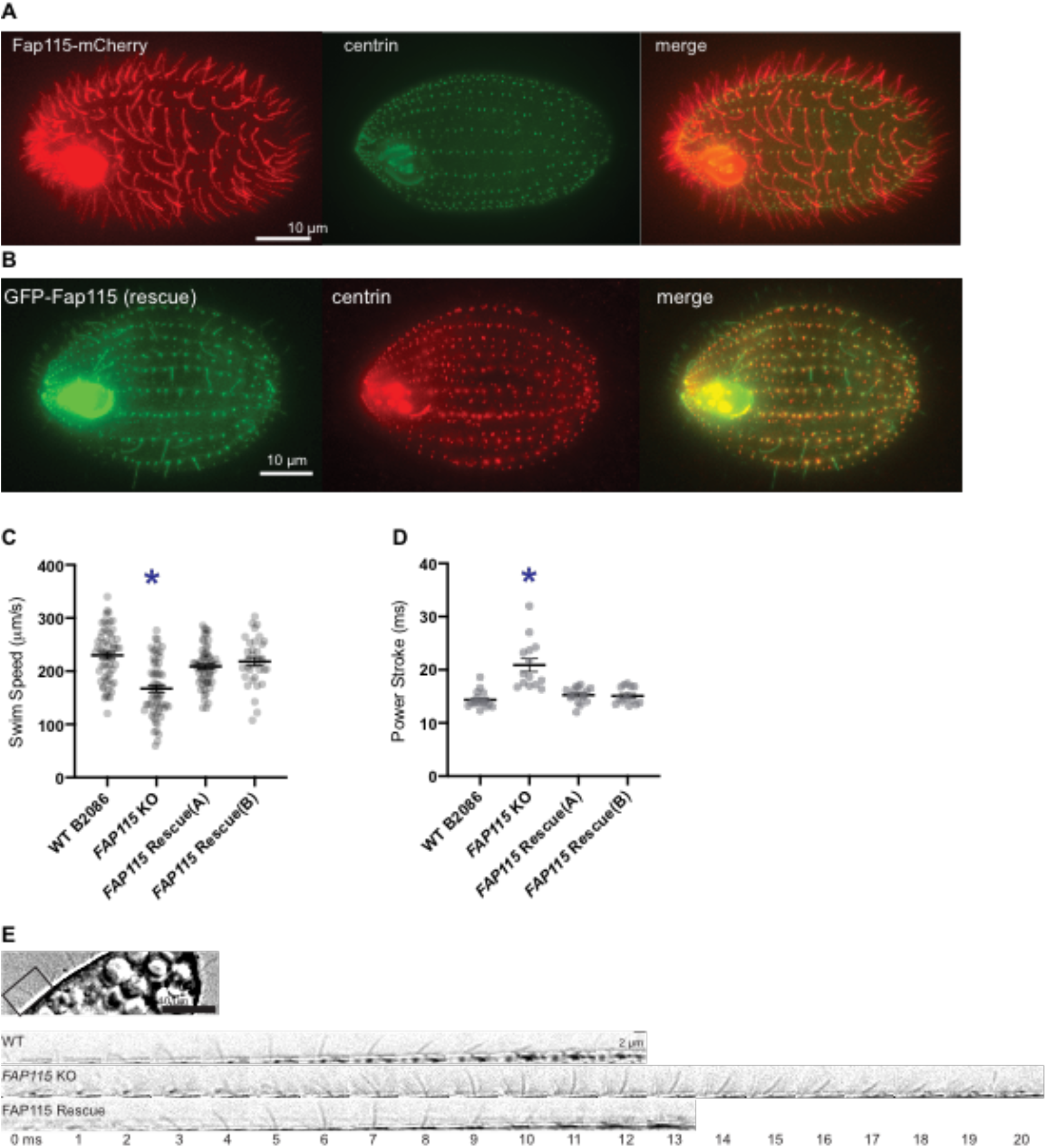
A) Fap115-mCherry localizes to cilia and colocalizes with the basal body protein centrin as detected by antibody staining (see Materials and Methods). B) GFP-Fap115 localizes to cilia and basal bodies in the *FAP115* KO C) Swim speed measurements in wild-type (WT), *FAP115* KO, and *FAP115* rescue strains D) Duration of individual cilia power strokes measured in control, *FAP115* KO, and *FAP115* rescue strains based on images such as those in E) sample image of a *Tetrahymena* cell, showing identification of a single cilium (top) and single frames from time-lapse images of cilia power strokes measured in WT, *FAP115* KO, and *FAP115* rescue strains (bottom). *p<0.01, error bars=SEM

Cilia and flagella microtubules must be built to resist mechanical strain during beating (Portran et al., 2017; Xu et al., 2017). We know that MIPs play a role in stability of the axoneme because loss of Rib72 renders axonemes flexible during the typically rigid force generating power stroke (Stoddard et al., 2018). Additionally, loss of MIPs in *Chlamydomonas* axonemes caused a stark decrease in structural stability of doublet microtubules (Owa et al., 2019).

*Tetrahymena FAP115* KO cells showed a decreased swim speed when compared to wild-type cells, which is restored to a normal speed with the reintroduction of Fap115 to the *FAP115* KO *Tetrahymena* strain (GFP-Fap115 rescue, Fig. 2B, C). To assess the possible cause of the decreased swim speed, we analyzed the ciliary density of the *FAP115* KO. Ciliary density was not altered (Fig. S2A), so we performed high-speed imaging of the ciliary beat pattern. High-speed time-lapse images of free-swimming cells were recorded, and movement of individual cilia was followed. A defect in the shape, frequency, or coordination of ciliary beating could affect the speed of swimming. There was no clear deformation of the cilia during the power stroke, as is seen when Rib72 is lost, but the duration of the power stroke was elongated in the knockout (Fig. 2D,E). While significant, the swim speed and power stroke defects were not as pronounced as the defects previously seen in *RIB72A, RIB72B*, or *RIB72A/B* double KO cells (Stoddard et al., 2018). *FAP115* KO *Tetrahymena* cells did not differ from wild type in growth rate or phagocytosis (Fig. S2B,C).

It stands to reason that either A-tubule MIPs besides Fap115 are individually necessary for stabilization of the axoneme or that the collective connection of A-tubule MIPs works together to stabilize the axoneme. Much like we see when analyzing ciliary beating defects, it would be useful in the future to determine whether there are specific axoneme-stabilizing MIPs or whether the stability breaks down suddenly after loss of a number of MIPs. This would suggest a concerted function for the stability of cilia. This finding does show that there are gradations of defects caused by losing individual MIPs and the whole system is not broken by the loss of a single MIP. In future studies, it will be important to determine the exact contributions of each MIP or a particular subset of MIPs to ciliary beating efficiency.

### Fap115 localizes to the A-tubule lumen (MIP6a)

We confirmed the localization *of Tetrahymena* Fap115 by performing Cryo-ET and subtomorgram averaging on isolated axonemes. *FAP115* KO axonemes showed complete loss of a single bi-lobed MIP density in the A tubule connecting protofilament (pf) A02 and A03 that has 8nm longitudinal periodicity in the WT axonemes (Fig. 3A-E). The loss of a single density in the *FAP115* KO, as compared with loss of multiple MIP densities in the *RIB72A* or *RIB72B* KO axonemes (Stoddard et al., 2018), is consistent with the more subtle ciliary phenotypes seen in the *FAP115* KO. These experiments demonstrate that Fap115 localizes to pf A02-03 in the lumen of the A-tubule, presumably interacting with Rib72A/B. The loss of the Fap115 density was also observed previously in the *RIB72A* or *RIB72B* KO (Stoddard et al., 2018), suggesting the dependence of Rib72A/B for the recruitment of Fap115 at pf A02 and A03 during the cilia assembly.

**Figure 3.**
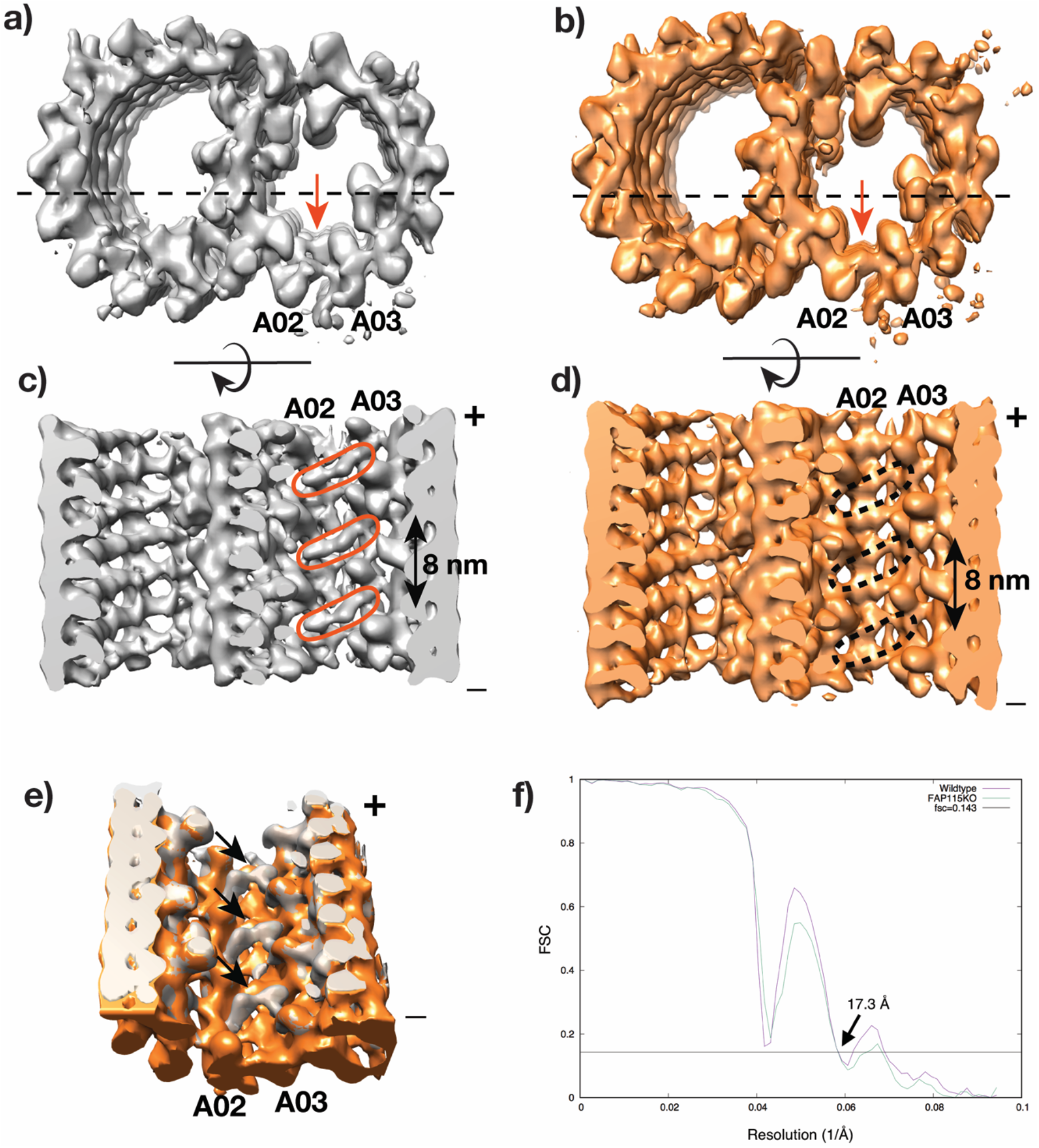
CryoET and subtomogram average of the axoneme doublet from the wild type and *FAP115* KO. a) and b) are the longitudinal view of the averages. Wild-type (grey) and *FAP115* KO (orange) are viewed from the minus ends of the doublet. The red arrows indicate the missing density in the KO axoneme that is present in the wild type. The dashed lines indicate the cross-section direction in c) and d). In c), the red elliptical circles highlight the density likely attributed to Fap115, the equivalent positions in the mutant doublet in d), where the density is missing, are indicated by black dash circle. e) is the overlay of two averages. The black arrows indicate the likely positions of Fap115 in the wild-type structure. They show 8 nm periodicity. f) Both structures are at 17.3 Å, determined by the FSC at 0.143 cutoff.

Across phylogeny, the presence of MIP homologs in motile cilia-containing organisms suggests high conservation of MIP structural architecture. In the two systems with the most highly characterized MIPs, *Chlamydomonas* and *Tetrahymena*, the localization and globular densities of MIPs appear to be highly similar (Ichikawa et al., 2017; Ma et al., 2019, Khalifa 2020). Dependence of Fap115 localization on Rib72A/B is consistent with high-resolution structures in *Chlamydomonas* showing interactions between Rib72 and Fap115 (Ma et al., 2019). It should be noted that Fap115 in *Tetrahymena* is four times the size of Fap115 in *Chlamydomonas*. Because both proteins are made up of multiple EF-hand domains, it is conceivable that these proteins function and localize in a similar manner. It is also possible that other systems have multiple proteins that share the role of a single MIP in *Tetrahymena*.

Much of the work on stability of the axoneme will be intimately tied to how axonemes are assembled and the timing of MIP integration into the axoneme. Axoneme microtubules assemble an order of magnitude slower than cytoplasmic microtubules and it is probable that MIPs are being incorporated into doublet microtubules coincident with tubulin addition to the axoneme (Sánchez and Dynlacht, 2016). The absence of Fap115 in the Rib72 knockout cells suggests a sequential and hierarchical assembly of MIPs. Rib72A/B are still present in the *FAP115* KO, but Fap115, along with most other A-tubule MIPs, is absent in *RIB72A/B* KOs. There is evidence that local curvature of the doublet microtubules is affected by MIP structures (Ichikawa et al., 2017, 2019; Khalifa et al., 2020), but specifically how the curvature is created and how it may affect recruitment and binding of additional MIPs is yet to be understood. Structural specificity of microtubules and direct binding to tubulin or other MIPs are likely mechanisms for early assembly of the MIP architectures.

Many questions about MIPs remain, and future research in the field is necessary to flush out the identities, structures, and functions of the lumenal meshwork of proteins inside of microtubules.

## Materials and Methods

### *Tetrahymena* Strains

The wild-type (B2086.2) *Tetrahymena* strain was obtained from the *Tetrahymena* Stock Center (Cornell University, Ithica, NY). All strains were grown in the standard medium 2% SPP at 30°C. RIB72A/B single and double KO strains are described in Stoddard et al., 2018.

Endogenous macronuclear *FAP115* (Ttherm_00193760) in B2086.2 cells was tagged with a C-terminal mCherry from the 4T2-1 vector containing C-terminal *FAP115* sequence and downstream sequence as homology for recombination. The *FAP115* macronuclear knockout was created by replacing the macronuclear *FAP115* gene in B2086.2 with the fragment of 4T2-1, containing mCherry and paromomycin resistance, using 0.5-1 kb homology from upstream and downstream regions of the gene. *Tetrahymena* were transformed with particle bombardment (Cassidy-Hanley et al., 1997; Hai and Gorovsky, 1997), then transformants were identified by resistance to paromomycin. For the *FAP115* KO, paromomycin concentrations were increased until the cells were fully assorted as knockouts, confirmed with RT-PCR. We used 2 independently assorted *FAP115* KO strains in our analyses.

To create an exogenous expression rescue strain, *FAP115* cDNA amplified from wildtype *Tetrahymena* cells was cloned into a plasmid containing RPL29 homology arms (pBS-MTT-GFP-gtw; see Stoddard et al., 2018). The construct was linearized and introduced to *FAP115* KO *Tetrahymena* cells by biolistic bombardment. Two independently assorted cycloheximide resistant strains were saved and used for analysis. Induction using CdCl2 (0.5 μg/ml) was between 1 hour and 24 hours, as noted.

### Proteomics

*Tetrahymena* ciliary axonemes were isolated from wild-type (B2086.2), single knockout *(RIB72A* or *RIB72B*), and double knockout strains (*RIB72A/RIB72B*) via calcium shock (Rosenbaum and Carlson, 1969). These axonemes were pelleted by centrifugation then solubilized, reduced and alkylated with 4% (w/v) sodium dodecyl sulfate, 10mM Tris (2-carboxyethyl) phosphine hydrochloride (TCEP), 40mM chloroacetamide, 100mM Tris pH 8.5 and boiled at 95°C for 10 minutes. Axonemes were then sonicated with a probe sonicator (Branson) for 1 second on, then 1 second off for a total of 1 minute. Approximately 100 micrograms of total protein were processed using the filter-aided sample preparation (FASP) method (Wiśniewski, 2016). Briefly, samples were diluted 10-fold with 8M Urea, 0.1M Tris pH8.5 and applied to the top of an Amicon Ultra 0.5mL 30kD NMWL cutoff (Millipore) ultrafiltration device. Samples were washed in the devices three time with 8M Urea, 0.1M Tris pH8.5, and another three times with 2M Urea, 0.1M Tris pH8.5. Endoproteinase Lys-C (Wako) was added first at a 1:100 protease to protein ratio then incubated approximately 2 hours on a nutator at room temperature. Trypsin (Pierce) was add next also at 1:100 then incubated overnight again on a nutator at room temperature. Tryptic peptides were eluted via centrifugation and desalted using a C-18 spin column (Pierce) according to the manufacture instructions. Samples were suspended in 3% (v/v) acetonitrile/0.1% (v/v) trifluoroacetic acid for 0.5 μg/uL peptide by A280nm and 1uL was directly injected onto a C18 1.7 μm, 130 Å, 75 μm X 250 mm M-class column (Waters), using a Waters M-class UPLC. Peptides were gradient eluted at 300 nL/minute from 3% to 20% acetonitrile over 100 min into an Orbitrap Fusion mass spectrometer (Thermo Scientific). Precursor mass spectra (MS1) were acquired at a resolution of 120,000 from 380 to 1500 m/z with an AGC target of 2E5 and a maximum injection time of 50 ms. Dynamic exclusion was set for 20 s with a mass tolerance of +/− 1Ó ppm. Precursor peptide ion isolation width for MS2 fragment scans was set at 1.6 Da using the quadrupole, and the most intense ions were sequenced by Top Speed with a 3 second cycle time. All MS2 sequencing was performed using higher energy collision dissociation (HCD) at 35% collision energy and scanned out in the linear ion trap. An AGC target of 1E4 and 35 second maximum injection time was used. Orbitrap data rawfiles were searched against the Uniprot *Tetrahymena thermophila* database (05-04-2020) using Maxquant with cysteine carbamidomethylation as a fixed modification and methionine oxidation and protein N-terminal acetylation as variable modifications. All peptides and proteins were thresholded at a 1% false discovery rate (FDR).

### Homology assessment of Rib72A/B and Fap115

Analysis of MIP conservation was performed using existing genetic data from a diverse taxonomic sample composed of 29 different model organisms (21 with motile cilia and 8 without). Taxon sampling was adapted from previous phylogenetic analysis of microtubule-based organelles (Carvalho-Santos et al., 2010, 2011). Only model organisms with accessible whole genome sequences were considered for analysis to be confident about homology assessment. Amino acid sequences for Rib72A/B and Fap115 were taken from the *Tetrahymena* Genome Database (Ciliate.org). Protein homology was determined by BLASTp search across the non-redundant protein sequence database (nr), including the respective model organism identifier in the search set using the BLASTp algorithm with default parameters. A reciprocal best hit BLAST search was done for the top search result in each species to determine whether the protein of interest is also the best hit in the *T. thermophila* genome and to discriminate between orthologues and paralogues (Bork et al., 1998; Tatusov et al., 1997). To do this, the highest scoring accession was then searched in BLAST using the same settings but modifying the search set to only include *T. thermophila* as the organism. If the protein of interest was the best hit in the reciprocal search, then it was considered homologous. Proteins in the taxon sampling were confirmed to be homologous if the protein BLAST yielded a result with an E-value less than or equal to 0.001 and had a positive reciprocal best hit BLAST to the protein of interest in *T. thermophila*.

### Fluorescence microscopy

Cells were fixed and imaged as described previously (Stoddard et al., 2018). Briefly, log phase cells were pelleted, then resuspended in a 3% formaldehyde solution. Cells were incubated in primary α-TtCen1 (Alexander J. Stemm-Wolf et al., 2005) antibody at 4°C overnight and secondary antibody (Alexa 488 or 594 anti-rabbit) for 1h at room temperature. Antibodies were diluted 1:1000 in PBS + 1% bovine serum albumen (BSA). Cells were washed three times with PBS + 0.1% BSA after each antibody incubation. Cells were mounted in Citifluor (EMS, Hatfield, PA) for viewing. Projection images were generated using ImageJ software.

### Swim speed assay

*Tetrahymena* cells were grown at 30°C in SPP media (2% protease peptone, 0.1% yeast extract, 0.2% glucose, 0.003% FeCl3) to mid–log phase, 2×10^5^ cells/ml. Cells were then placed on a slide and imaged using differential interference contrast (DIC) imaging with a 20× objective at 10 frames/s on a Nikon Ti microscope (Nikon, Japan) using NIS Elements software (Nikon). ImageJ with the ImageJ software plugin MTrackJ (E. Meijering) was used to track and quantify the movement of *Tetrahymena* cells. Twenty cells were quantified per condition, and the experiment was performed in triplicate.

### Cilia wave beating analysis

Log-phase *Tetrahymena* cells were place in a drop on cover glass, then DIC images of free-swimming cells (above the plane of the cover glass) were captured at 996 frames/sec with a 100x NA1.45 oil immersion objective on a Nikon Ti-E microscope (Nikon, Japan) with a Hamamatsu Flash4.0 V3 camera fitted with a V3 Firebird Camlink Board, using Nikon Elements software. Images were analyzed using ImageJ.

### Cell growth assay

Wild-type and *FAP115* KO *Tetrahymena* cells were grown at 30°C in 2%SPP medium to a concentration of 2 x 10^4^ cells/mL, as measured by a hemocytometer. Cell density was quantified by hemocytometer every two hours for eight hours, then at 24 hours. This experiment was completed in duplicate.

### Phagocytosis assay

*Tetrahymena* cells were incubated with 0.5% India ink in 1% SPP media at 25°C. At 30 minutes after the India ink was added, cells were washed in 2% SPP and fixed with 2% formaldehyde. Cells were mounted on slides and the average number of food vacuoles per cell was quantified (*n* =20 cells) using phase contrast imaging with a 10× objective on an upright light microscope (Nikon Ti Japan). This experiment was completed in duplicate.

### Cryo sample preparation and cryo-ET

*Chlamydomonas* fap115 ko cryo-ET and Image processing were performed as reported previously (Stoddard et al., 2018).

For cryoET on *Tetrahymena* axonemes, 10 nm colloid gold (Sigma) was mixed with ciliary axonemes isolated from the wild-type or *FAP115* KO. 4 μl sample was applied onto Quantifoil grid Rh/Cu 200 R2/2 (Quantifoil, Inc), and was flash frozen into liquid ethane using a Vitrobot (FEI, Inc).

Tomography tilt series were collected on a field emission gun (FEG) microscope (Titan Krios, FEI, Inc) operated at 300kV. The microscope was equipped with a post-column energy filter Bio-Quantum GIF (Gatan, Inc). The slit width was set at 25 eV. SerialEM (Mastronarde, 2005) was used for the tomography tilt series collection. Images were recorded on a K3 direct electron detector (Gatan, Inc) in the super-resolution and dose-fractionation mode. The nominal magnification is 33,000, the effective physical pixel size is 2.65 Å. The specimen was tilted in 2° increment in two-branch, starting from zero degree, in a range of −60° to +60°. The total accumulative dose on the sample was limited to 80 electron/Å^2^.

For tomogram reconstruction and subtomogram averaging, the dose-fractionated movie at each tilt in the tilt series was corrected of motion and summed using MotionCor2 (Zheng, et al 2017). The tilt series were aligned based on the gold bead fiducials by use IMOD (Kremer et al., 1996) and TomoAlign (Fernandez et al., 2018). The contrast transfer function for each tilt series was determined and corrected by TomoCTF (Fernandez, 2006). The tomograms were reconstructed by TomoRec (Fernandez et al., 2019) that took into account of the beam-induced sample motion during data collection. Respectively, for the wild-type, 5235 subtomograms from 15 datasets, for the *FAP115* KO construct, 5926 subtomograms from 18 datasets were used for the subtomogram averaging. The subtomogram alignment and average were carried out, without using any external reference, by the program MLTOMO implemented in the Xmipp software package (Scheres et al., 2009). Extensive classification of subtomogram was carried out in order to find the 16nm periodicity of the Rib72A/B. The subtomograms were shifted accordingly in the longitudinal direction of the axoneme so that these MIPs are in-register. This was followed by additional alignment in a “gold-standard” scheme in 2x-binned format (effective pixel size 5.30 Å) (Scheres and Chen, 2012). The final resolution, based on the Fourier Shell Correlation (FSC) cutoff at 0.143, is reported as 17.3 Å for both the wild-type and the *FAP115* KO structure (Fig. 3F). The structures have been deposited in EMDB with access codes EMD-22712 (B2086, WT) and EMD-22713 (*FAP115* KO).

## Acknowledgements

This work was supported by funding from the National Institutes of Health (NIH) R01GM127571 (PI - M.W.); R35GM118099 (PI – D.A.). Cryo-ET sample preparation was performed by Fei Guo at the UC Davis BioEM facility. The UC Davis BioEM Facility is supported by the user fees, the Department of Molecular and Cellular Biology, the College of Biosciences, the Office of Research and the Provost’s Office. The Technical Director, Dr. Fei Guo, is supported by discretionary funds provided by Professor Jodi Nunnari (MCB). Cryo-ET data were collected at the Louise Mashal Gabbay Cellular Visualization Facility at Brandeis University (Waltham, MA) and at UCSF. We thank David Bulkley and Glenn Gilbert in the Bay Area CryoEM Center at UCSF for technical support. This center is supported in part by NIH grants including 1S10OD020054 and 1S10OD021741. We also thank Daryn Baker and Steve Hinds for maintaining computational infrastructure at SCU to allow for assessment of homology. We thank Rachel Howard-Till and Usha Kar for critical review.

---

**Table S1:**
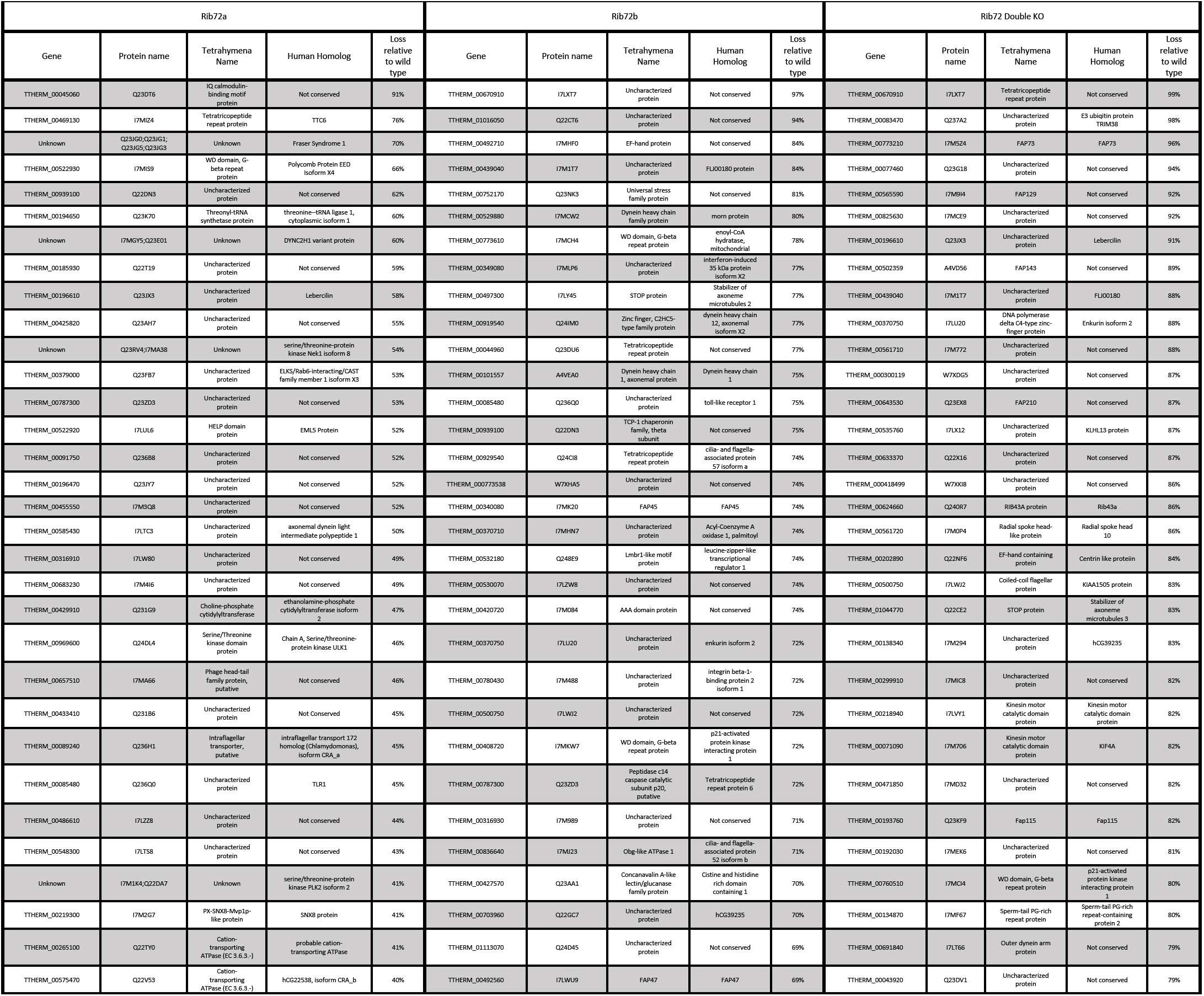
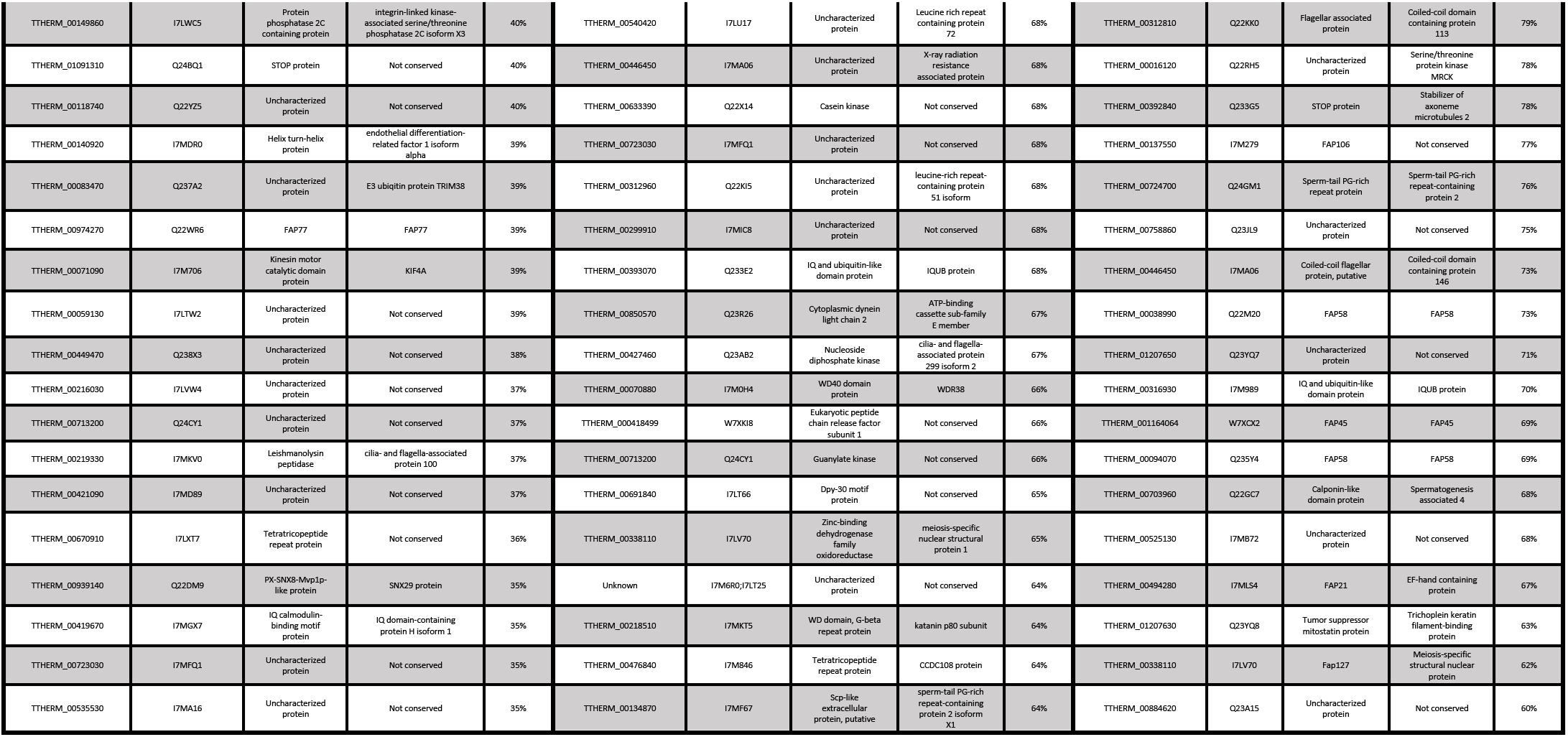
Top 50 proteins lost in *RIB72AB* KO, *RIB72A* KO, or *RIB72B* KO cells

**Figure S1.**
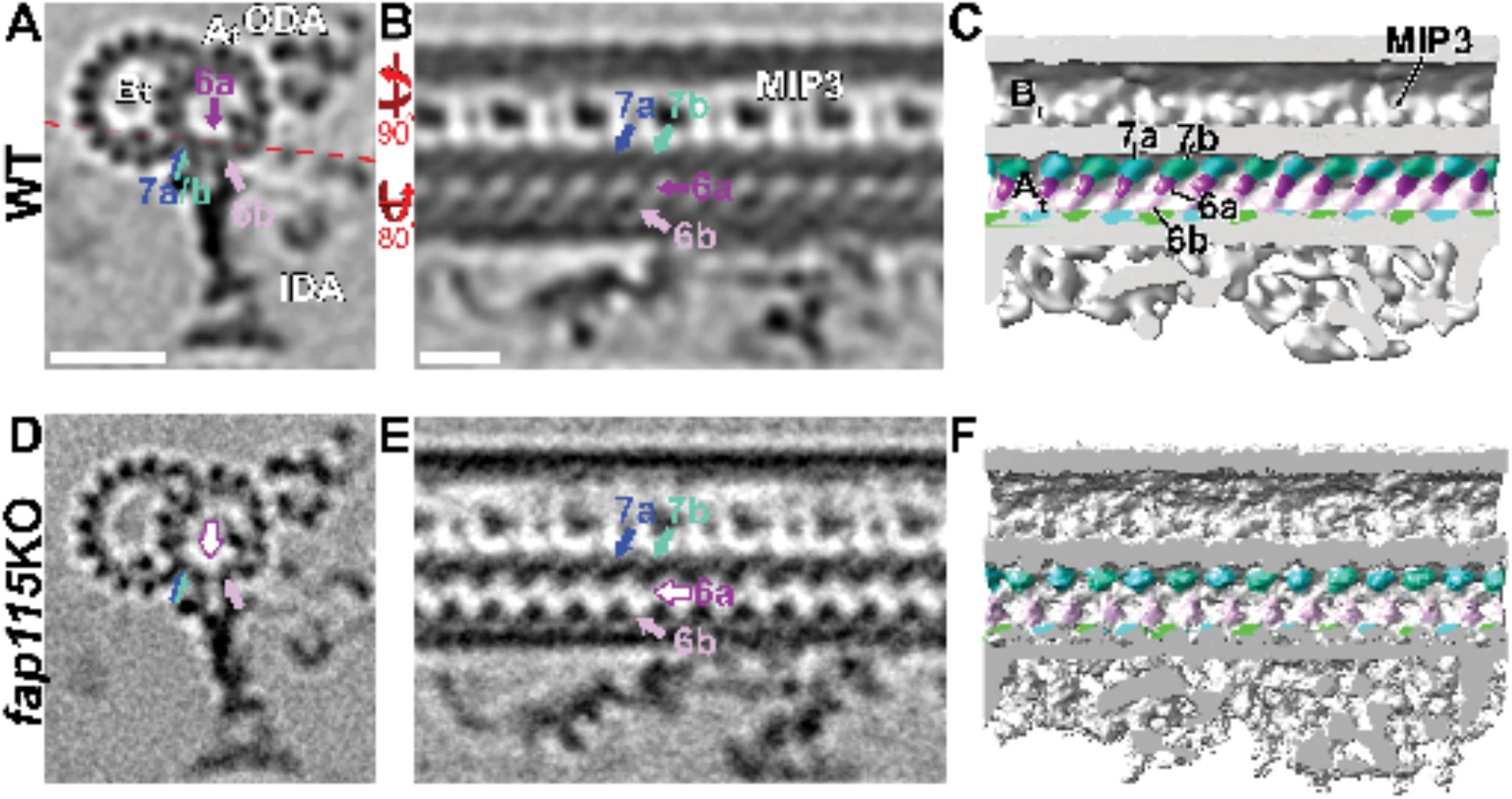
*Chlamydomonas* - CryoET and subtomogram average of the axoneme doublet from the wild type and fap115 KO mutant

**Figure S2.**
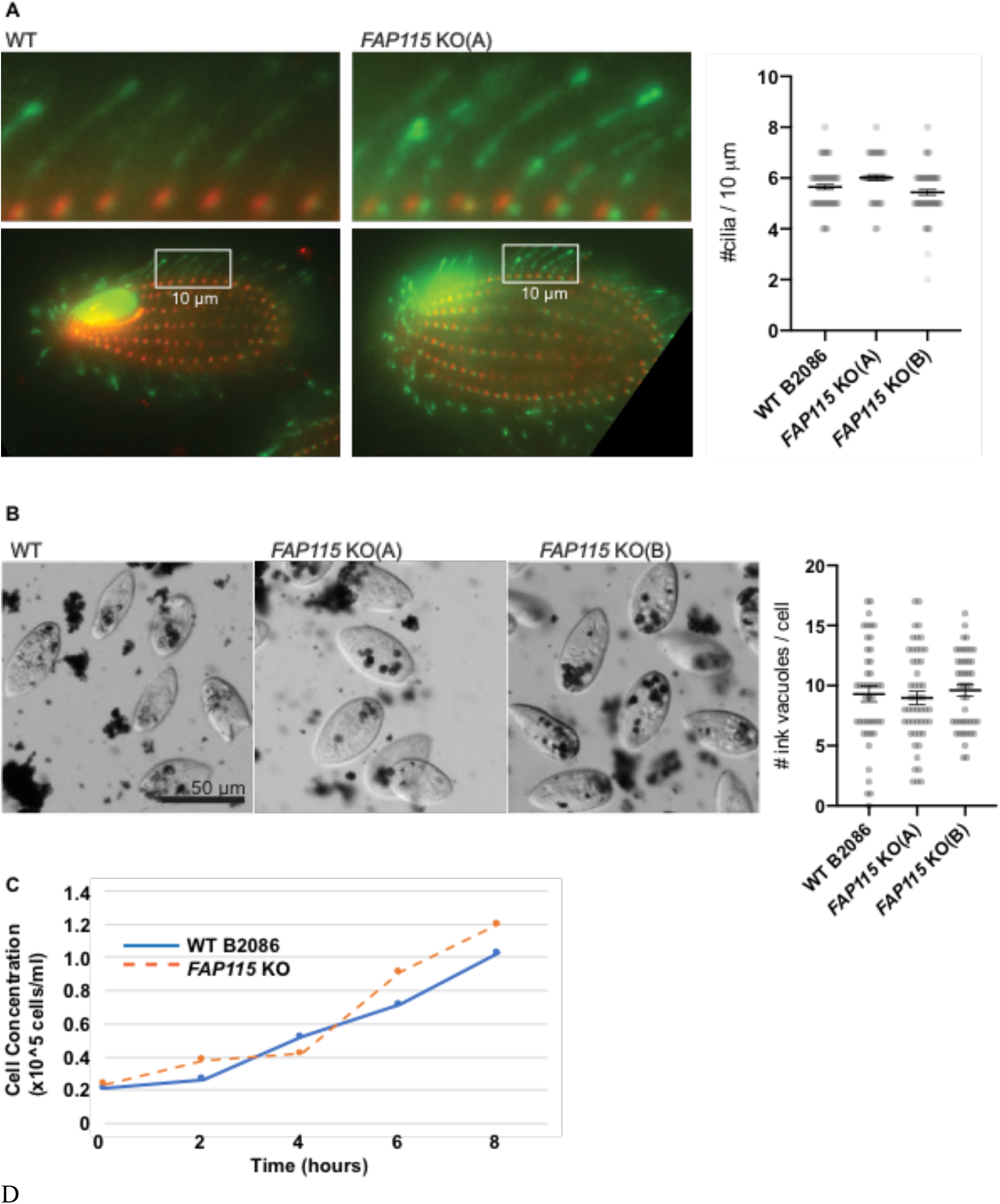
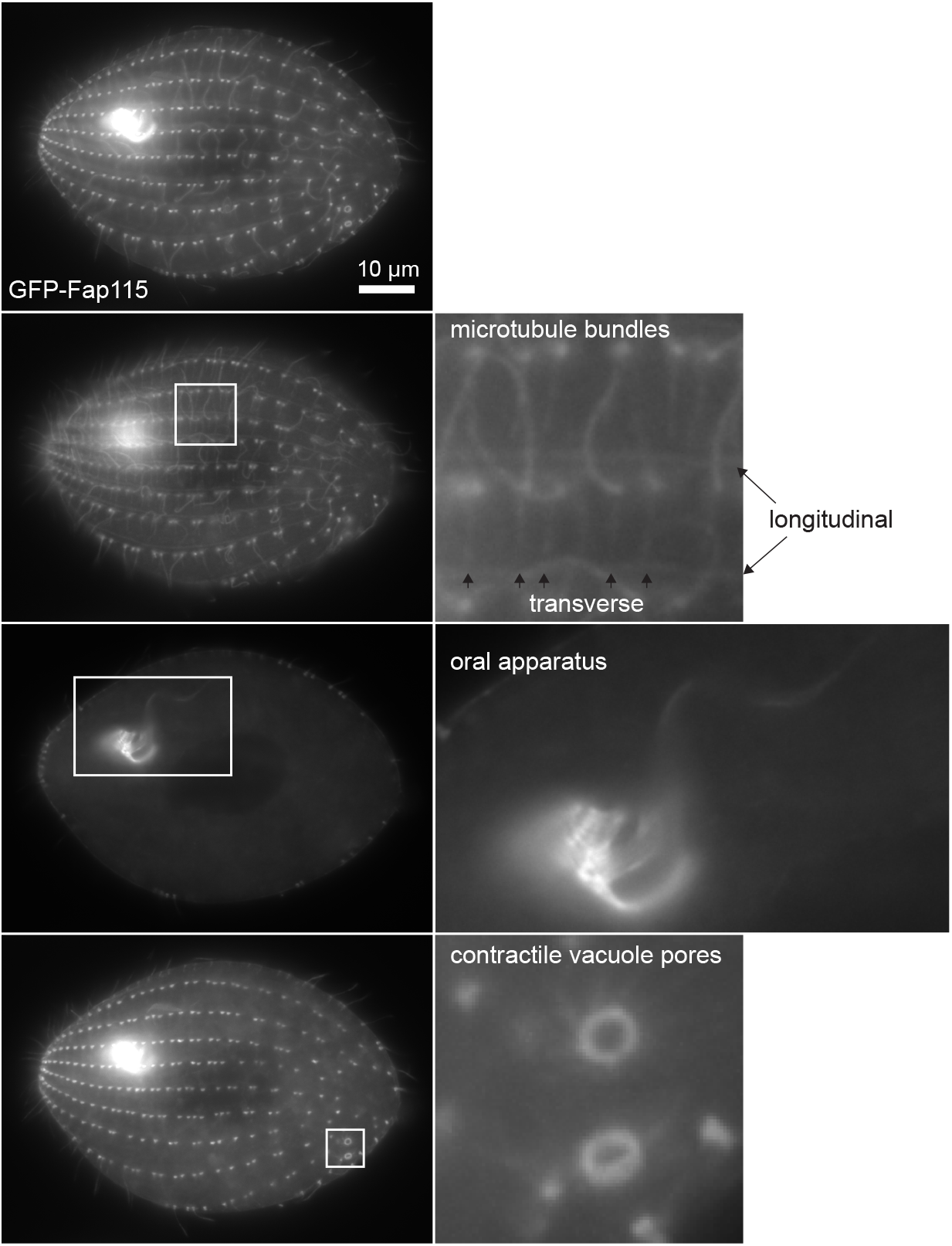
Additional phenotypic analysis *FAP115* KO and Rescue cells. A) Ink vacuoles from a phagocytosis assay are present after exposure to ink in media. Chart shows number of ink vacuoles counted in control and *FAP115* KO strains. B) Cilia density within a 10μm region in the center of the cell. Boxed region from bottom panel shown above. Chart shows cilia density in WT and *FAP115* KO strains. C) Growth curves for WT and *FAP115* KO strains. D) GFP-Fap115 localizes to additional microtubule-based structures when overexpressed in the *FAP115* KO strain.

